# Normative Brain Size Variation and the Remodeling of Brain Shape in Humans

**DOI:** 10.1101/205930

**Authors:** P. K. Reardon, Simon N. Vandekar, Siyuan Liu, Raihaan Patel, Min Tae M. Park, Aaron Alexander-Bloch, Jakob Seidlitz, Liv S. Clasen, Jonathan D. Blumenthal, Jay N. Giedd, Ruben C. Gur, Raquel E. Gur, Jason P. Lerch, M. Mallar Chakravarty, Theodore D. Satterthwaite, Russel T. Shinohara, Armin Raznahan

## Abstract

Evolutionary and developmental increases in primate brain size have been accompanied by systematic shifts in the proportionality of different primate brain systems. However, it remains unknown if and how brain patterning varies across the more than 2-fold inter-individual variation in brain size that occurs amongst typically-developing humans. Using *in vivo* neuroimaging data from 2 independent cohorts totaling nearly 3000 individuals, we find that larger-brained humans show preferential areal expansion within specific fronto-parietal cortical networks (default mode, dorsal attentional) and related subcortical regions, at the expense of primary sensory/motor systems. This targeted areal expansion recapitulates cortical remodeling across evolution, manifests by early childhood and is linked to molecular signatures of heightened metabolic cost. Our results define a new organizing principle in human brain patterning which governs the highly-coordinated remodeling of human brain shape as a function of naturally-occurring variations in brain size.

**One Sentence Summary:** A hodologically and metabolically expensive brain network is preferentially expanded in larger-brained humans.

## Main Text

Total brain size can vary over 2-fold amongst typically developing humans of the same age, and this variability is greater still when considering clinical populations (*1*). Brain size variation has been robustly linked to coordinated changes in the proportional size of different brain systems across mammalian evolution and development (*2–4*), but it remains unknown if analogous organizational shifts accompany the striking inter-individual brain-size variation that exists in humans. While there is emerging evidence that normative brain size variation within humans is accompanied by a systematic change in the proportional volume of individual structures such as the cerebellum and thalamus (*5, 6*), we still lack a spatially comprehensive, fine-grained map of size-dependent anatomical reorganization of the human brain. Such an anatomical scaling map would illuminate the differential organizational requirements of larger vs. smaller brains, and represent a fundamental advance in our basic knowledge of human brain patterning in health.

Here, we leverage two large-scale *in vivo* structural neuroimaging datasets to provide the first high-resolution anatomical atlas of areal scaling across the human brain. We discover topographically complex patterns of non-linear areal scaling across the human cortex, which give rise to systematic differences in the organization of cortical anatomy in smaller vs. larger brained individuals. We then use integrative analysis across diverse *in vivo* and *post mortem* assays to show that these interregional differences in cortical scaling are aligned with inter-regional differences in: cortical expansion during evolution and development (*2*), functional connectivity (*7*), gene expression (*8*), and resource consumption (*9*). Finally, by extending our analysis beyond the cortex, we reveal analogous patterns of non-linear scaling in the human subcortex, which lead to the highly-coordinated remodeling of subcortical shape as a function of inter-individual variations in thalamic, pallidal, striatal, amygdalar and hippocampal size. Taken together, these cortical and subcortical findings detail a general shift in the balance between associative and sensorimotor brain systems as a function of naturally-occurring brain size variation in humans. The scaling relationships which govern this anatomical reorganization recapitulate those that have been shown to operate across evolutionary and developmental time scales, and provide an empirical bridge between macroscopic (shape) and microscopic (molecular) features of the human brain.

Our study harnesses 2,904 structural magnetic resonance images (sMRI) brain scans from two independent cohorts – (i) a discovery National Institutes of Health (NIH) sample of 1531 longitudinally acquired brain scans from a 1.5 Tesla MRI machine in 792 youth aged 5-25 years (338 females, **Table S1A, Fig S1A**, (*10*)), and (ii) a replication Philadelphia Neurodevelopmental Cohort (PNC) sample of 1,373 crosssectional scans from a 3 Tesla MRI machine in youth aged 8-23 years (728 females, **Table S1B, Fig S1B**, (*11*)). In both datasets, we first used well-validated tools for automated shape analysis from sMRI to measure local surface area at a total of ~80k points across the cortical sheet (*10*). To generate a reference map of areal scaling in the cortex, we adapted an analytic method for the study of scaling initially proposed by Galileo Galilei (*12*) and since refined in phylogenetic research (*13*). Thus, we modelled inter-individual differences in the surface area of each cortical vertex as a function of inter-individual differences in total cortical surface area within a log-log regression framework. The slopes of these regressions quantified the scaling relationship between area at each vertex and total cortical surface area, such that a scaling coefficient of 1 at every vertex would represent the null hypothesis that areal patterning of the cortical sheet is stable across differences in cortical size (i.e. no changes in proportional vertex areas with cortical size). In this framework, scaling coefficients greater than 1 represent supra-linear or “positive scaling” (i.e. vertex area becomes disproportionately large with increasing cortical size), while coefficients less than 1 represent sub-linear or “negative scaling” (i.e. vertex area becomes disproportionately small with increasing cortical size). These log-log regressions were implemented within semiparametric regression models that simultaneously accounted for both linear and nonlinear age and sex effects on surface area at each vertex (**Methods**, **Table S2**, (*14*)).

Analyses in the discovery NIH sample revealed that scaling relationships between local surface area and total surface area vary widely across the human cortex, and result in a differential patterning of the cortical mantle whereby certain regions become relatively larger with increasing cortical size (e.g. medial prefrontal) and other regions become relatively smaller (e.g. primary visual) (Fig 1A). These spatial gradients in areal scaling show (i) strong bilateral symmetry, (ii) stability between ages 5 and 25 years, and (iii) equivalence in males and females (**Methods**). Next, by testing for a statistically-significant deviation of the scaling coefficient at each vertex from 1 (i.e. the null hypothesis of linear scaling), and then correcting for multiple comparisons (**Methods**), we defined two distributed cortical domains that display statistically-significant deviations from linear scaling (i.e. regions of “positive scaling” and “negative scaling”), and are therefore robustly re-shaped as a function of inter-individual differences in total cortical size (Fig 1B). Our findings in the NIH sample were highly reproducible in the independent PNC replication dataset - both at the level of continuous scaling gradients (Fig 1C), and domains of statistically significant non-linear scaling that survive correction for multiple comparisons (Fig 1D). We were able to quantify the statistical significance of this visible agreement between NIH and PNC scaling maps using a novel surface-based permutation method (*15*) which generates an empirical null overlap distribution by repeatedly “spinning” one map against the other using a well-established spherical projection of the cortical sheet (*10*) (p < 0.001 for observed spatial overlap vs. empirical null distribution, **Methods**). Collectively, these convergent results in two large independent human datasets firmly establish that the larger cortical sheets of bigger-brained individuals show disproportionate areal expansion of dorso-lateral prefrontal, lateral temporo-parietal, and medial parietal multimodal cortices, alongside a relative areal contraction of unimodal cortices surrounding the calcarine (visual) and central (somatosensory and motor) sulci. We next harnessed spatial permutation testing to directly compare the cortical scaling map in our discovery NIH cohort (Fig 1B) to several other key features of cortical organization from independent studies.

**Figure 1.**
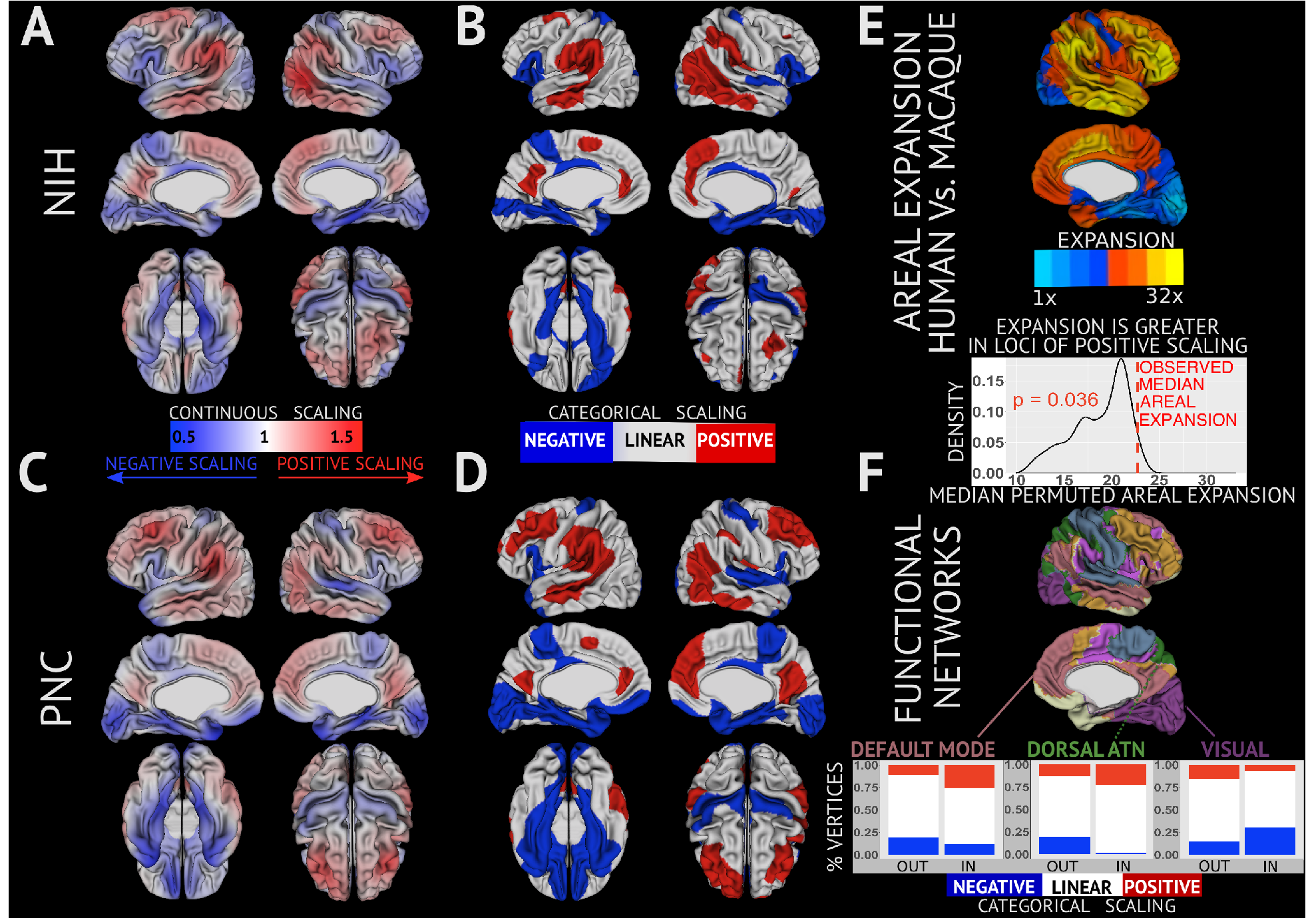
Maps of areal scaling in the human cortex and links with cortical evolution and function. Continuous **(A)**, and categorical **(B)** maps of areal scaling in the NIH cohort. The continuous scaling map **(A)** shows un-thresholded coefficients for individual vertex area scaling with total cortical surface area. Color hue denotes the direction of deviation from linear scaling (red - positive scaling, blue- negative scaling), and greater color intensity denotes a greater magnitude of deviation from linear scaling. The categorical map **(B)** uses block colors to identify vertices with statistically-significant positive (red) or negative (blue) deviations from linear scaling. Continuous **(C)**, and categorical **(D)** maps of areal scaling in the PNC cohort. **E)** An independently published vertex-level map of areal expansion in humans vs. macaques, with a density plot showing that the observed inter-species areal expansion score in human positive scaling regions is significantly elevated relative to a null distribution generated by one-thousand spatial permutations of the NIH categorical scaling map. **F)** An independently-published map of resting state functional connectivity networks in the human brain (*7*), with a bar plot illustrating the significant non-random overlap between 3 resting-state networks (each p < 0.001). and regions of negative vs. positive scaling in the NIH sample.

First - to test if cortical reorganization as a function of brain size variation in humans recapitulates known patterns of cortical reorganization with brain size variation across primates - we assessed the convergence between domains of significant positive and negative areal scaling in the NIH sample and a previously published cortical map of relative areal expansion/contraction in humans vs. macaques (Fig 1E) (*2*). Visually, both forms of brain size variation appear to be associated within similar domains of positive (frontal and lateral temporal cortices) and negative areal scaling (primary sensori-motor and medial temporal cortices). Spatial permutation testing confirmed this visual impression, and established that cortical domains of positive areal scaling within humans display a statistically-significant (p = 0.036) degree of overlap with domains of cortical expansion in humans relative to macaques (and other simian primates (*4*)), which have, in turn, been shown to be disproportionately expanded in adult vs. infant humans (*2*). These results indicate that possession of a larger brain - either through evolution, development, or standing inter-individual size variation - is accompanied by the disproportionate focal areal expansion of prefrontal, lateral temporo-parietal and medial parietal regions, alongside the relative areal contraction of primary visual and motor cortices.

We hypothesized that such a highly-conserved pattern of differential growth investment between domains of positive vs. negative cortical scaling should show some relationship with the functional topography of the cortical sheet (*16*). To test this hypothesis, we directly co-registered the categorical NIH cortical scaling map (Fig 1B) with a canonical parcellation of the cortical sheet into 7 large-scale functional networks defined by patterns of resting-state functional connectivity (Fig 1F) (*4*), and tested each network for spatial correspondence with cortical domains of relative areal expansion or contraction. Spatial permutation testing identified three resting-state networks that showed a statistically-significant spatial association with cortical scaling: (i) the “default mode” network (DMN, p < 0.001, enriched for domains of positive scaling), (ii) the dorsal attentional network (DAN, p < 0.001, enriched for positive scaling), and (iii) the visual network (p < 0.001, enriched for negative scaling (**Methods**). The DMN and DAN are two canonical distributed associative systems (*17–20*) that display a preferential involvement in highly-abstracted functions spanning social cognition, prospection, memory, valuation, moral decision making and executive control (*21, 22*). These functional specializations are thought to be supported by a conspicuously large number of costly long-range connections (*23, 24*) with other brain networks (*25*), which endow the DMN and DAN with a special capacity to integrate information across multiple subnetworks in the brain (*26*). In contrast, the visual network –– which scales sub-linearly in size with the cortical sheet –– is known to be relatively isolated in terms of its connectivity with other brain networks (*27*). Thus, size-dependent shifts in areal organization of the human cortex are partly aligned with the modular organization of functional connectivity between cortical regions, such that increasing brain size is accompanied by a redistribution of surface area from lower-order sensory cortices to integrative associative systems.

We next sought to probe molecular signatures that might be associated with the regionally-specific investments in cortical growth identified by our scaling analyses. To achieve this objective, we aligned the categorical NIH cortical scaling map with a publically available atlas of gene expression from six adult human brains (Allen Institute for Brain Sciences, AIBS, (*8*)), and ranked ~16k genes detailed within the AIBS atlas by their differential expression between cortical regions of significant positive vs. negative scaling (Fig 2A, **Methods**). Gene ontology (GO) enrichment analyses of this ranked gene list (**Methods**, (*28*)) revealed that high-ranking genes (i.e. those with greater expression in regions of positive vs. negative scaling) were significantly associated with: specific cellular components - mitochondrion and the post-synaptic membrane; and specific cellular processes - respiratory electron transport chain assembly/function and K+ channel activity (Fig 2B, **Table S3A-C**). The upregulation of energetic pathways in cortical domains with positive scaling was robustly replicated using an independent bioinformatics approach (*29*) involving GO enrichment analysis of gene sets showing statistically significant differential expression regions of positive vs. negative scaling (**Methods, Table S4A-C**). These analyses hint that cortical regions which undergo preferential areal expansion with increasing cortical size may also be the recipients of a preferential energetic investment - as reflected by elevated transcriptional activity in mitochondrial pathways for oxidative phosphorylation. To probe this energetic hypothesis using an orthogonal data modality, we compared the NIH cortical scaling to an independently-published cortical map (*30*) of resting-state arterial blood flow as estimated through imaging of arterial spin labelling (ASL) in the PNC cohort (**Methods**, Fig 2C). Inter-regional differences in resting state arterial blood flow provide a well-established proxy for inter-regional differences in resting state oxygen consumption (*31*), allowing us to ask if domains of positive areal scaling show a conspicuously high consumption of oxygenated blood at rest relative to domains of negative areal scaling. A formal test of this hypothesis using spatial permutation (**Methods**) confirmed that regions of positive scaling in the NIH map do indeed show elevated perfusion at rest relative to regions of negative scaling (p = 0.028). Therefore, both gene-expression and blood flow data support the inference that cortical regions of relative areal expansion with increasing brain size amongst humans are distinguished from other cortical regions by a heighted energetic cost.

**Figure 2.**
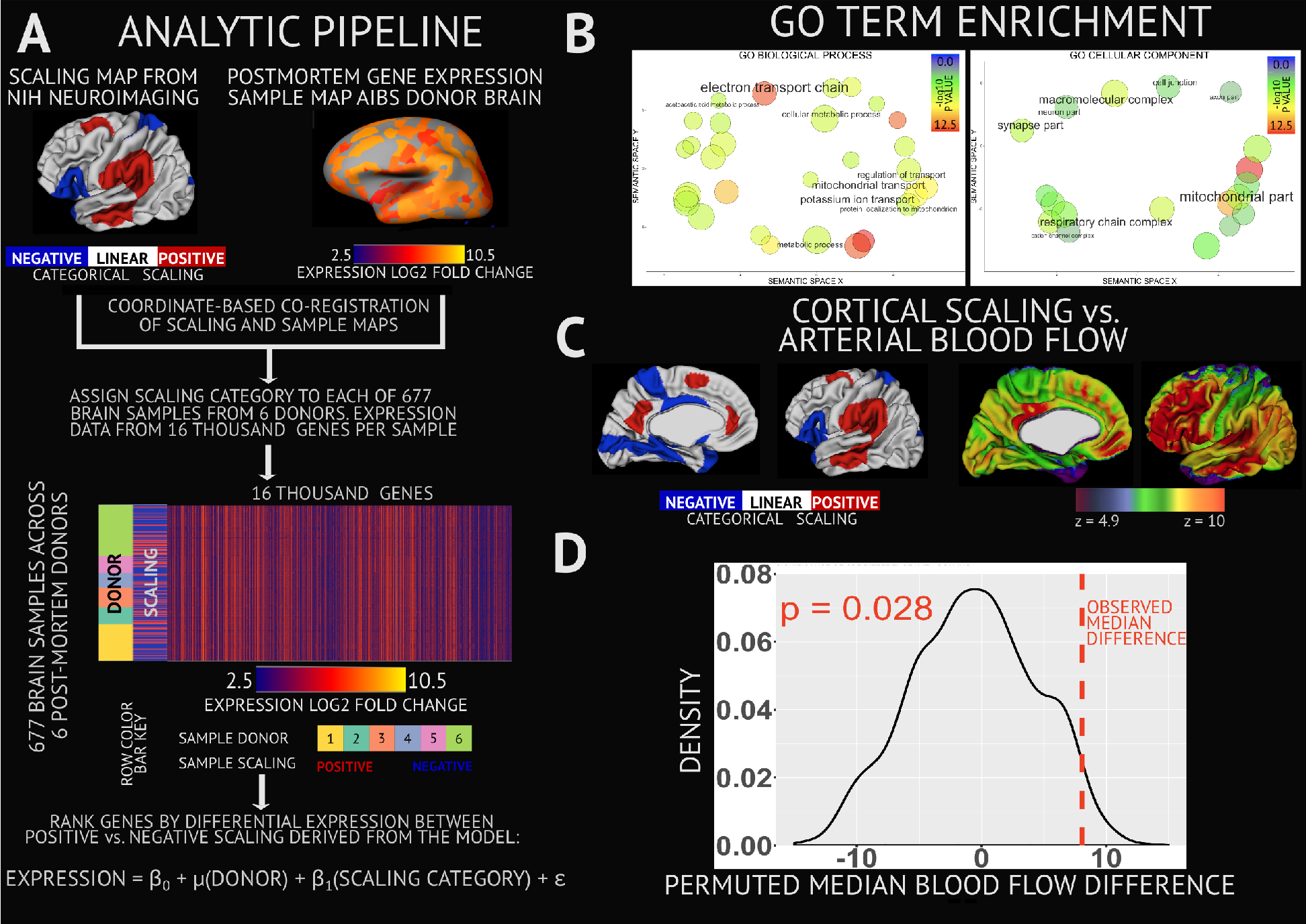
Linking Cortical Scaling to Cortical Gene Expression and Metabolism. **A)** Cartoon of analytic pipeline for integration of cortical scaling maps with postmortem microarray gene expression data from the 6 donor brains in the Allen Institute of Brain Sciences (AIBS). This pipeline ranks ~16k genes by the difference in their expression value between regions of positive vs. negative scaling. **B)** Gene Ontology (GO) term enrichment analysis of this ranked gene list indicates that regions of positive scaling show elevated relative expression of genes coding for mitochondrial and synaptically-located proteins with roles in oxidative metabolism and K^+^ signaling (respectively). Significantly enriched GO terms are plotted in semantic space such that similar terms are represented close to one another. Circles are scaled and colored according to the −log_10_ of the *p*-value for the significance of each term. (**C)** NIH scaling map next to projection of Arterial Spin Labelling (ASL) resting state blood flow map from Satterthwaite et al (*27*) - with warmer colors indicating higher blood flow. A spatial permutation test established that observed differences in median blood flow between regions of positive vs. negative scaling is significantly (p = 0.028) greater than chance.

Collectively, these analyses of areal scaling and its correlates advance our understanding of cortical organization in a number of key directions. First, we establish
that the relative expansion of associative vs. sensorimotor cortices with increasing cortical size is not limited to primate evolution and development (*2, 4*), but is also seen along the third major axis of brain size variation - inter-individual differences in size within a species. The consistent emergence of this scaling map strongly suggests that a shared mechanism is operating to link regional and global cortical surface areas in three very different contexts: inter-species size variation across evolutionary time-frames, intra-specific size variation across developmental time, and intra-specific size variation across individuals. We reason that this shared mechanism must involve at least one of two processes: (i) the laying down of regional differences in cortical cell count and/or cell size during prenatal corticogenesis (*32, 33*), and (ii) the subsequent emergence of regional differences in cellular morphology and/or neuropil composition (*34, 35*). These two candidate mechanisms predict distinct histological differences between cortical regions of positive vs. negative scaling - yielding specific falsifiable hypothesis which are increasingly tractable given improving access to suitable postmortem tissue from humans (https://neurobiobank.nih.gov) and non-human primates (e.g. http://www.chimpanzeebrain.org) of varying ages and statures.

Second, because brain growth is achieved at a metabolic cost (*36*), regional differences in areal scaling signal disparate biological investment between different cortical regions as a function of increasing brain size. Our analyses of transcriptomic and hemodynamic data reinforce this theme of greater biological investment in regions of positive vs. negative areal scaling (Fig 2). The tight conservation of this scaling map across phylogenetic (*2, 4*), ontogenic (*2, 37*) and inter-individual (current report) forms of brain size variation further suggests that disproportionate expansion of prefrontal, medial parietal and lateral parieto-temporal cortices is a fundamental and adaptive organizational feature of larger vs. smaller cortices. A candidate functional basis for this recurrent organizational motif is provided by our observation that domains of positive areal scaling tend to encompass the DMN and DAN. These brain networks are topologically specialized for the integration of information across multiple lower-order brain regions from numerous long range connections (*17–20*). We speculate that the positive areal scaling of DMN and DAN cortices may reflect or facilitate (*38, 39*) the computational requirements associated with their specialized integrative role in the brain. More specifically, computational complexity theory predicts that many integrative algorithms will experience supra-linear increases in computational load with a linear increase in the size of its inputs (*40*), and we observe a supra-linear increase in the physical size of integrative cortices (DMN and DAN) given a linear increase in the size of those input cortices that they integrate information across (e.g. sensorimotor systems).

The highly organized and conserved remodeling of cortical structures as a function of inter-individual differences in brain size suggests that a similar phenomenon may operate for subcortical systems with which the cortex is densely inter-connected (*41*). As a first step towards addressing this question, and determining if the cortex is distinctive in the complexity of its size-dependent reorganization, we generalized our analytic approach (**Methods**) to each of five different subcortical structures: thalamus, pallidum, striatum, amygdala, and hippocampus. Specifically, for each structure, we estimated how the surface area of individual structure vertices scales with total structure surface area. For each of the subcortical structure examined, we identified topographically complex scaling patterns that were bilaterally symmetric, and broadly organized along medio-lateral and rostro-caudal gradients. Further echoing our cortical observations, patterns of non-linear areal scaling across the surface of each subcortical structure were stable across the age-ranges examined, equivalent between males and females, and highly reproducible across the NIH and PNC cohorts (Fig 3). Testing for statistically-significant deviation of scaling coefficients from 1, and correcting for multiple comparisons, defined distributed facets of each structure showing robust changes in relative surface area as a function of total structure size (Fig 3 B,D). Domains of statistically-significant positive and negative scaling were highly coordinated within and between subcortical structures such that proximate facets of different structures tended to show identical deviations from linear scaling. Consequently, we detect an extended domain of positive subcortical areal scaling which encompasses the rostral amygdala, overlying regions of the hippocampal head plus adjacent dorso-lateral facets of the hippocampal tail, and predominantly dorsal facets of the caudate, putamen and thalamus. In contrast, domains of negative scaling tended to encompass ventral and/or medial facets of the hippocampus, putamen and thalamus.

**Figure 3.**
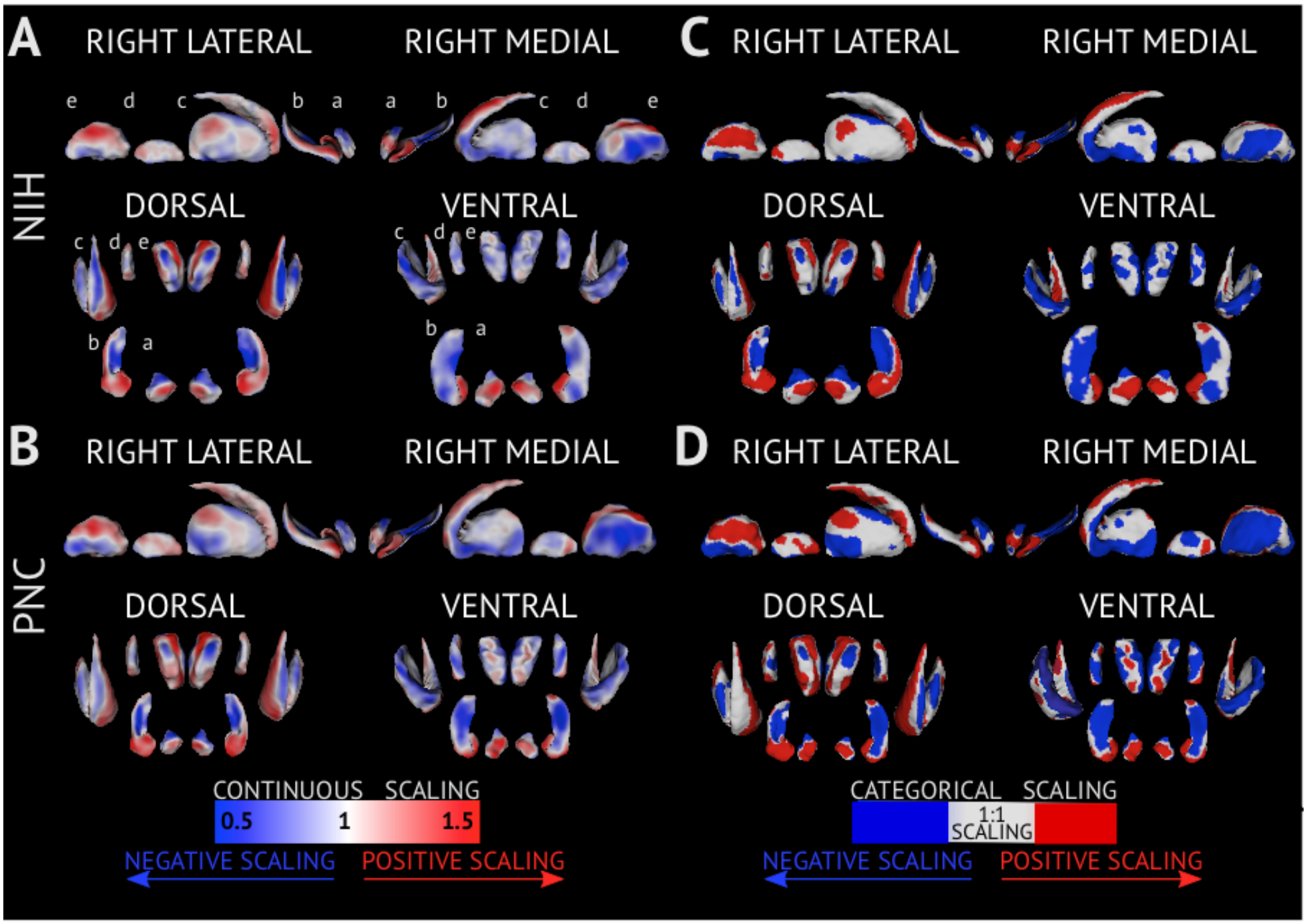
Maps of areal scaling in the human subcortex. Each panel provides views of the amygdala (a), hippocampus (b), striatum (c), globus pallidus (d) and thalamus (e) from four different anatomical vantage points. **A, B)** Vertex-level **(A)**, and categorical **(B)** maps of areal scaling in the NIH cohort. The continuous scaling map **(A)** shows un-thresholded coefficients for scaling of area at individual vertices with the area of their parent structure. Color hue denotes the direction of deviation from linear scaling (red - positive scaling, blue-negative scaling), and greater color intensity denotes a greater magnitude of deviation from linear scaling. The categorical map **(B)** uses block colors to identify vertices with statistically-significant positive (red) or negative (blue) deviations from linear scaling. **C, D)** Vertex-level **(C)**, and categorical **(D)** maps of areal scaling in the PNC cohort.

Strikingly, in many instances the observed spatial patterning of subcortical scaling recapitulated that of the cortex according to known topographies of cortical-subcortical connectivity. For example, medial and ventro-lateral facets of the thalamus that become disproportionately small with increasing thalamic size are known to show preferential functional connectivity to visual, limbic and motor domains of negative areal scaling in the cortex, while dorsomedial domains of positive thalamic areal scaling show preferential connectivity with the DMN (*42*). Similarly coordinated patterns of subcortio-cortical scaling were apparent in the striatum (*43*), and amygdalo-hippocampus (*44*). Thus, analysis of areal scaling in the subcortex demonstrates that the cortex is not alone in undergoing marked size-dependent remodeling, and reinforces the trend that larger brained humans show preferential expansion of associative vs. sensorimotor systems.

Taken together our analyses of anatomical scaling within the human cortex and subcortex provide reference maps that chart the profound reorganization of regional brain anatomy that accompanies naturally-occurring size variation amongst human brains. Tangential scaling gradients in the cortex recapitulate those that arise with evolutionary and developmental changes in brain size, and are closely coordinated with topographies of functional connectivity, gene expression and blood flow. The integration of our scaling maps with these orthogonal datasets highlights the DMN, DAN and linked subcortical regions as a distributed and integrative set of brain regions which are conspicuously “costly”, and become increasingly favored recipients of anabolic resources as brain size increases. Such insights recommend the analysis of macroanatomical scaling as a classical approach to the study of biological shape (*12*) that is ripe for revival in modern neuroscience.

## ACKNOWLEDGMENTS

This research was supported in part by the Intramural Research Program of the NIMH (Clinical trial reg. no. NCT00001246, clinicaltrials.gov; NIH Annual Report Number, ZIA MH002794, Protocol ID 89-M-0006).

## REFERENCES

1. J. N. Giedd et al., Brain development during childhood and adolescence: a longitudinal MRI study. Nat. Neurosci. 2, 861–863 (1999).

2. J. Hill et al., Similar patterns of cortical expansion during human development and evolution. Proc. Natl. Acad. Sci. U. S. A. 107, 13135–40 (2010).

3. L. Krubitzer, The magnificent compromise: cortical field evolution in mammals. Neuron. 56, 201–208 (2007).

4. T. A. Chaplin, H.-H. Yu, J. G. M. Soares, R. Gattass, M. G. P. Rosa, A conserved pattern of differential expansion of cortical areas in simian primates. J. Neurosci. Off. J. Soc. Neurosci. 33, 15120–15125 (2013).

5. C. Mankiw et al., Allometric Analysis Detects Brain Size-Independent Effects of Sex and Sex Chromosome Complement on Human Cerebellar Organization. J. Neurosci. Off. J. Soc. Neurosci. 37, 5221–5231 (2017).

6. P. K. Reardon et al., An Allometric Analysis of Sex and Sex Chromosome Dosage Effects on Subcortical Anatomy in Humans. J. Neurosci. Off. J. Soc. Neurosci. 36, 2438–2448 (2016).

7. B. T. T. Yeo et al., The organization of the human cerebral cortex estimated by intrinsic functional connectivity. J. Neurophysiol. 106, 1125–1165 (2011).

8. M. J. Hawrylycz et al., An anatomically comprehensive atlas of the adult human brain transcriptome. Nature. 489, 391–9 (2012).

9. S. N. Vaishnavi et al., Regional aerobic glycolysis in the human brain. Proc. Natl. Acad.Sci. U. S. A. 107, 17757–17762 (2010).

10. J. N. Giedd et al., Child psychiatry branch of the national institute of mental health longitudinal structural magnetic resonance imaging study of human brain development. Neuropsychopharmacology. 40, 43–9 (2015).

11. T. D. Satterthwaite et al., Neuroimaging of the Philadelphia neurodevelopmental cohort. Neuroimage. 86, 544–53 (2014).

12. G. Galilei, Dialogues Concerning Two New Sciences (Macmillan, New York, 1914).

13. S. J. Gould, Allometry and size in ontogeny and phylogeny. Biol. Rev. Camb. Philos. Soc. 41, 587–640 (1966).

14. S. Wood, Generalized additive models: an introduction with R (CRC press, 2006).

15. S. N. Vandekar et al., Topologically dissociable patterns of development of the human cerebral cortex. J. Neurosci. Off. J. Soc. Neurosci. 35, 599–609 (2015).

16. A. Raznahan et al., Patterns of Coordinated Anatomical Change in Human Cortical Development: A Longitudinal Neuroimaging Study of Maturational Coupling. Neuron. 72, 873–884 (2011).

17. P. Hagmann et al., Mapping the Structural Core of Human Cerebral Cortex. PLOS Biol. 6, e159 (2008).

18. N. A. Crossley et al., Cognitive relevance of the community structure of the human brain functional coactivation network. Proc. Natl. Acad. Sci. 110, 11583–11588 (2013).

19. S. M. Smith et al., Correspondence of the brain’s functional architecture during activation and rest. Proc. Natl. Acad. Sci. 106, 13040–13045 (2009).

20. F. de Pasquale et al., A cortical core for dynamic integration of functional networks in the resting human brain. Neuron. 74, 753–764 (2012).

21. M. E. Raichle, The brain’s default mode network. Annu. Rev. Neurosci. 38, 433–447 (2015).

22. D. S. Margulies et al., Situating the default-mode network along a principal gradient of macroscale cortical organization. Proc. Natl. Acad. Sci. U. S. A. 113, 12574–12579 (2016).

23. A. F. Alexander-Bloch et al., The anatomical distance of functional connections predicts brain network topology in health and schizophrenia. Cereb Cortex. 23, 127–38 (2013).

24. M. P. van den Heuvel, O. Sporns, Rich-club organization of the human connectome. J. Neurosci. Off. J. Soc. Neurosci. 31, 15775–15786 (2011).

25. E. Bullmore, O. Sporns, The economy of brain network organization. Nat. Rev. Neurosci. 13, 336–349 (2012).

26. O. Sporns, Network attributes for segregation and integration in the human brain. Curr. Opin. Neurobiol. 23, 162–171 (2013).

27. B. T. T. Yeo, F. M. Krienen, M. W. L. Chee, R. L. Buckner, Estimates of segregation and overlap of functional connectivity networks in the human cerebral cortex. NeuroImage. 88, 212–227 (2014).

28. E. Eden, R. Navon, I. Steinfeld, D. Lipson, Z. Yakhini, GOrilla: a tool for discovery and visualization of enriched GO terms in ranked gene lists. BMC Bioinformatics. 10, 48 (2009).

29. M. V. Kuleshov et al., Enrichr: a comprehensive gene set enrichment analysis web server 2016 update. Nucleic Acids Res. 44, W90–97 (2016).

30. T. D. Satterthwaite et al., Impact of puberty on the evolution of cerebral perfusion during adolescence. Proc. Natl. Acad. Sci. 111, 8643–8648 (2014).

31. M. E. Raichle et al., A default mode of brain function. Proc. Natl. Acad. Sci. U. S. A. 98, 676–82 (2001).

32. I. Reillo, C. de Juan Romero, M. Á. Gartía-Cabezas, V. Borrell, A role for intermediate radial glia in the tangential expansion of the mammalian cerebral cortex. Cereb. Cortex N. Y. N 1991. 21, 1674–1694 (2011).

33. D. J. Cahalane, C. J. Charvet, B. L. Finlay, Systematic, balancing gradients in neuron density and number across the primate isocortex. Front. Neuroanat. 6, 28 (2012).

34. G. N. Elston, T. Oga, I. Fujita, Spinogenesis and pruning scales across functional hierarchies. J. Neurosci. Off. J. Soc. Neurosci. 29, 3271–3275 (2009).

35. G. N. Elston, R. Benavides-Piccione, J. DeFelipe, J. Neurosci. Off. J. Soc. Neurosci., in press.

36. C. W. Kuzawa et al., Metabolic costs and evolutionary implications of human brain development. Proc. Natl. Acad. Sci. 111, 13010–13015 (2014).

37. A. E. Lyall et al., Dynamic Development of Regional Cortical Thickness and Surface Area in Early Childhood. Cereb. Cortex N. Y. N 1991. 25, 2204–2212 (2015).

38. M. Rubinov, Constraints and spandrels of interareal connectomes. Nat. Commun. 7, 13812 (2016).

39. R. L. Buckner, F. M. Krienen, The evolution of distributed association networks in the human brain. Trends Cogn. Sci. 17, 648–665 (2013).

40. M. Sipser, Introduction to the theory of computation (Thompson Course Technology, ed. 2nd, 2006).

41. S. N. Haber, Corticostriatal circuitry. Dialogues Clin. Neurosci. 18, 7–21 (2016).

42. K. Hwang, M. Bertolero, W. Liu, M. D’Esposito, The human thalamus is an integrative hub for functional brain networks. J. Neurosci., 67–17 (2017).

43. E. Y. Choi, B. T. T. Yeo, R. L. Buckner, The organization of the human striatum estimated by intrinsic functional connectivity. J. Neurophysiol. 108, 2242–2263 (2012).

44. J. L. Robinson et al., Neurofunctional topography of the human hippocampus. Hum. Brain Mapp. 36, 5018–5037 (2015).

